# Stimulus-driven cerebrospinal fluid dynamics is impaired in glaucoma patients

**DOI:** 10.1101/2025.01.15.633258

**Authors:** Ji Won Bang, Carlos Parra, Kevin Yu, Hyun Seo Lee, Gadi Wollstein, Joel S. Schuman, Kevin C. Chan

## Abstract

Cerebrospinal fluid (CSF) dynamics, driven by sensory stimulation-induced neuronal activity, is crucial for maintaining homeostasis and clearing metabolic waste. However, it remains unclear whether such CSF flow is impaired in age-related neurodegenerative diseases of the visual system. This study addresses this gap by examining CSF flow during visual stimulation in glaucoma patients and healthy older adults using functional magnetic resonance imaging. The findings reveal that in glaucoma, CSF inflow becomes decoupled from visually evoked blood-oxygenation-level-dependent (BOLD) response. Furthermore, stimulus-locked CSF patterns, characterized by decreases following stimulus onset and increases after offset, diminish as glaucoma severity worsens. Mediation analysis suggests that this flattened CSF pattern is driven by a flatter BOLD slope, resulting in a shallower CSF trough and a reduced rebound. These findings unveil a novel pathophysiological mechanism underlying disrupted stimulation-driven CSF dynamics in glaucoma and highlight potential *in vivo* biomarkers for monitoring CSF in the glaucomatous brain.

## Main

Glaucoma is an age-related neurodegenerative disease characterized by the gradual loss of retinal ganglion cells in the eye and disruptions along the brain’s visual pathways^1,2^. Despite extensive research, the underlying cause of the disease remains unknown. Currently, intraocular pressure is the only clinically modifiable risk factor, yet glaucomatous damage can progress even when intraocular pressure is well-controlled^3^. Consequently, glaucoma remains the leading cause of irreversible blindness worldwide.

Recent research highlights the vital role of cerebrospinal fluid (CSF) flow in clearing metabolic waste and maintaining the health of the eye and brain^4,5^. While CSF has been implicated in the pathogenesis of glaucoma, direct evidence is limited. Existing studies on CSF flow in glaucoma have primarily focused on the optic nerve’s subarachnoid space during resting-state conditions, documenting slowed CSF flow^6,7^. Although these studies provide insights into local CSF dynamics, it remains poorly understood whether glaucoma also involves impaired CSF dynamics at a systemic level. This gap in understanding is significant, as glaucoma not only affects the optic nerve but also extends more broadly into the intracranial system, even in the early stages of the disease^8^. Supporting this, glaucoma patients exhibit elevated levels of amyloid β and tau, proteins linked to neurodegenerative diseases, along the visual pathway, including the retina^9,10^, lateral geniculate nucleus, and primary visual cortex^11^. Additionally, GABA, an inhibitory neurotransmitter vulnerable to amyloid β and tau toxicity^12-16^ and known to facilitate waste clearance^17^, is reduced in the visual cortex of glaucoma patients^1^. Elevated levels of amyloid β and tau may result from impaired metabolic waste clearance in the brain, while reduced GABA may hinder glymphatic clearance^17^. These findings underscore the critical need to investigate CSF dynamics within the intracranial system in glaucoma patients.

Sensory stimulation-induced neuronal activity plays a crucial role in driving CSF flow^18^. While vision is central to daily activities, neurons in the visual cortex of both glaucoma patients and experimental animal models exhibit reduced firing rates^19^ and delayed responses^20^ to visual stimuli. This raises an imperative question: is CSF flow driven by visual activity compromised in glaucoma patients? To address this gap, we recruited glaucoma patients and healthy control subjects to identify impairments in CSF flow during visual stimulation in glaucoma patients. Specifically, we focused on the fourth ventricle, whose vertical shape allows for the detection of incoming CSF signals^18,21,22^, making it ideal for studying systemic CSF dynamics. Using functional magnetic resonance imaging (fMRI), we captured blood-oxygenation-level-dependent (BOLD) signals from the visual cortex while subjects viewed a high-contrast flickering checkerboard in a slow block design. CSF inflow signals were extracted from the lowest slice of the fMRI volume, positioned at the fourth ventricle, which is sensitive to upward CSF flow into the brain^22^.

Our findings revealed an inverse relationship between the BOLD response in the visual cortex and CSF inflow across subjects: an increase in the early BOLD signal was followed by a decrease in CSF inflow, while a subsequent decrease in BOLD signal was followed by a peak in CSF inflow. However, as glaucoma severity increased, this pattern weakened, showing reduced coupling between BOLD and CSF inflow and a diminished CSF rebound. This disrupted CSF inflow dynamics can be attributed to the impaired BOLD response, as confirmed by mediation analysis. As the BOLD slope flattens, its ability to suppress CSF inflow diminishes, leading to a shallower CSF trough. This elevated trough then restricts the subsequent rebound, resulting in a diminished CSF surge. In summary, our results demonstrate that in glaucoma, the visually triggered BOLD slope in the visual cortex is impaired, contributing to a flattened CSF flow pattern with shallower troughs and weaker rebounds. This study provides compelling evidence that CSF flow alterations in glaucoma extend beyond the optic nerve, reaching the fourth ventricle and likely affecting other intracranial CSF spaces and neural tissues as a result of the diminishing inflow. Additionally, our study introduces a non-invasive method for monitoring CSF flow in glaucoma patients, offering new insights into the broader neurological impact of the disease.

## Results

Fifty-two individuals diagnosed with primary glaucoma and twenty-seven healthy individuals of comparable age underwent clinical ophthalmic exams and MRI scans (see **Table 1**. for demographic and clinical characteristics). Individuals with glaucoma were assigned to either early or advanced stage based on the average visual field mean deviation (MD) of the left and right eyes combined (OU) (early stage: average OU MD > -6.0dB, advanced stage: average OU MD <-6.0dB)^1,23^.

**Table 1:** Demographic and clinical characteristics of glaucoma and healthy subjects. [pRNFL: peripapillary retinal nerve fiber layer, mGCIPL: macular ganglion cell-inner plexiform layer, NRR: neuroretinal rim, C/D: cup-to-disc ratio, OD: oculus dexter (right eye), OS: oculus sinister (left eye)]

|  | Healthy control<br>(n=27; mean<br>± S.E.M.) | Early glaucoma<br>(n=23; mean<br>± S.E.M.) | Advanced glaucoma<br>(n=29; mean<br>± S.E.M.) | Group effect (one-way ANOVA or Welch or Chi-squared test) |
| --- | --- | --- | --- | --- |
| Age (years) | 63.93 ± 1.47 | 65.17 ± 1.59 | 66.14 ± 1.33 | F(2,76)=0.614, P=0.544 |
| Sex | 12M, 15F | 7M, 16F | 16M, 13F | $\chi^2(2)=3.18$, P=0.204 |
| Visual field mean deviation OD (dB) | -1.45 ± 0.54 | -2.69 ± 0.61 | -17.32 ± 1.48 | F<sub>welch</sub>(2,45.80)=50.35, P<0.001 |
| Visual field mean deviation OS (dB) | -1.40 ± 0.36 | -2.48 ± 0.65 | -13.83 ± 1.62 | F<sub>welch</sub>(2, 40.92)=27.85, P<0.001 |
| Foveal threshold OD (dB) | 35.50 ± 0.41 | 35.14 ± 0.56 | 30.34 ± 1.07 | F<sub>welch</sub>(2, 43.32)=10.10, P<0.001 |
| Foveal threshold OS (dB) | 35.80 ± 0.53 | 35.05 ± 0.43 | 31.39 ± 0.86 | F<sub>welch</sub>(2, 43.19)=9.59, P<0.001 |
| Optic nerve head C/D OD | 0.43 ± 0.03 | 0.72 ± 0.03 | 0.76 ± 0.02 | F(2,71)=39.670, P<0.001 |
| Optic nerve head C/D OS | 0.41 ± 0.03 | 0.68 ± 0.04 | 0.78 ± 0.02 | F(2,71)=39.441, P<0.001 |
| pRNFL thickness OD (μm) | 89.09 ± 1.63 | 73.17 ± 2.30 | 61.45 ± 1.81 | F(2,72)=52.762, P<0.001 |
| pRNFL thickness OS (μm) | 90.26 ± 1.55 | 73.91 ± 2.35 | 63.03 ± 2.07 | F(2,72)=45.79, P<0.001 |
| mGCIPL thickness OD (μm) | 78.05 ± 1.31 | 66.24 ± 1.64 | 57.77 ± 1.91 | F<sub>welch</sub>(2,42.32)=41.48, P<0.001 |
| mGCIPL thickness OS (μm) | 77.90 ± 1.69 | 67.57 ± 1.56 | 58.38 ± 1.28 | F(2,64)=43.489, P<0.001 |
| NRR area OD (mm<sup>2</sup>) | 1.29 ± 0.06 | 0.85 ± 0.06 | 0.66 ± 0.05 | F(2,71)=32.384, P<0.001 |
| NRR area OS (mm<sup>2</sup>) | 1.34 ± 0.04 | 0.84 ± 0.07 | 0.65 ± 0.04 | F(2,71)=51.628, P<0.001 |

### Visual cortex BOLD response to stimuli is reduced in individuals with glaucoma

We initially evaluated whether our flickering checkerboard visual stimuli, presented in a slow block design (8s ON and 16s OFF, repeated 22 times across 2 runs) elicited diminishing BOLD signals from the visual cortex as glaucoma severity increased (**Fig. 1a,b**). Our results confirmed that the BOLD amplitude [11-12s post-stimulus onset; 0.72% for healthy controls, 0.57% for early stage, 0.41% increase for advanced stage; main effect of group, F_welch_(2, 45.390)=14.254, P<0.001, partial η^2^=0.301; healthy controls vs. early glaucoma, Games-Howell-corrected P=0.059, 95% confidence interval (CI)=-0.0045 to 0.2980; healthy controls vs. advanced glaucoma, Games-Howell-corrected P<0.001, 95% CI=0.1601 to 0.4521; early glaucoma vs. advanced glaucoma, Games-Howell-corrected P=0.004, 95% CI=0.0443 to 0.2743] and the percentage of voxels in the visual cortex activated by stimulation [46.71% for healthy controls, 43.21% for early stage, 34.02% for advanced stage; main effect of group, F(2,72)=16.842, P<0.001, partial η^2^=0.319; healthy controls vs. early glaucoma, Bonferroni-corrected P=0.443, 95% CI=-2.359 to 9.353; healthy controls vs. advanced glaucoma, Bonferroni-corrected P<0.001, 95% CI=7.106 to 18.271; early glaucoma vs. advanced glaucoma, Bonferroni-corrected P<0.001, 95% CI=3.544 to 14.839] declined with increasing glaucoma severity (**Fig. 1c-f**). This impact of glaucoma was not detected in the global brain BOLD signal, which represents the average BOLD signal across the entire gray matter mask (see **Extended Data** Fig. 1).

**Fig. 1:**
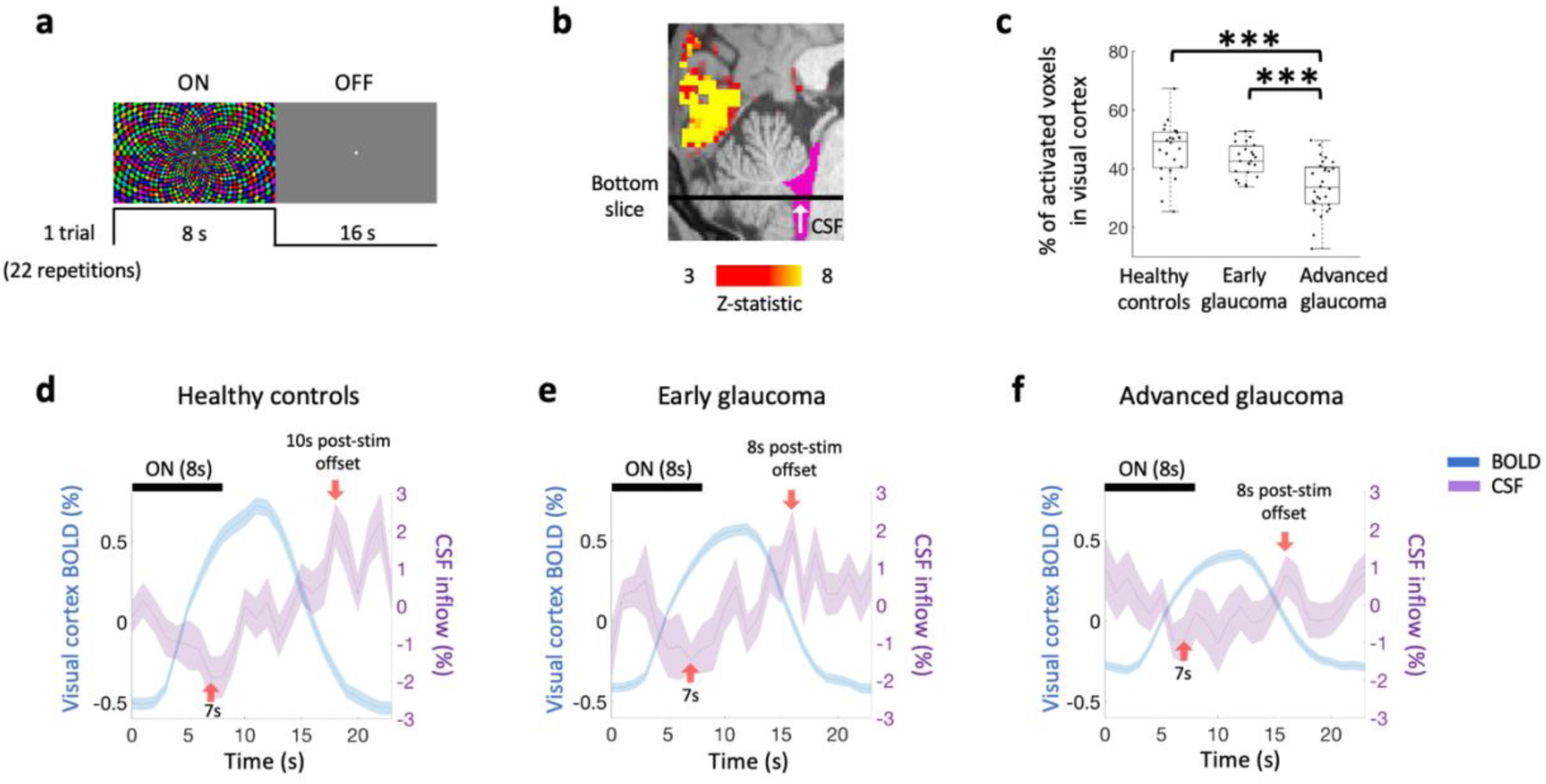
Stimulation-induced dynamics of cerebrospinal fluid degraded with glaucoma severity. **a**, Exemplar visual stimulus. A flickering checkerboard was presented for 8s (“On” condition), followed by a gray screen for 16s (“Off” condition) in each trial, repeated 22 times in total. The visual tasks consisted of 2 fMRI runs with 11 trials each. **b,** Exemplar position where the bottom axial imaging slice was positioned (black line) during the fMRI scan to measure incoming CSF inflow (white arrow) towards the fourth ventricle (pink). The visually evoked BOLD activation map (warm color) was overlaid on the sagittal anatomical MRI in the posterior brain. The color bar shows the z-values of activated voxels for the visual stimuli. **c,** The percentage of voxels in the visual cortex activated by stimulation was reduced with increasing glaucoma severity. The data are represented using box and whisker plots. In descending order, the lines in the plots represent: maximum, third quartile, median, first quartile, and minimum. ***Bonferroni-corrected P<0.001. **d-f,** The time courses of visual cortex BOLD signals (blue) and CSF (purple) in **d,** healthy controls, **e,** individuals with early glaucoma, and **f,** individuals with advanced glaucoma indicated that both the BOLD response and stimulus-locked CSF inflow were compromised in glaucoma. In d-f, the data represent mean ± S.E.M. across subjects. Red arrows indicate the CSF trough at 7s post-stimulus onset and the CSF surge at 8s or 10s post-stimulus offset.

### Stimulus-locked pattern of CSF inflow is attenuated in individuals with glaucoma

Under healthy conditions, bursts of neuronal activity induce arteriolar dilation, leading to an increased blood volume through neurovascular coupling^24^. This rise in blood volume displaces CSF to maintain constant intracranial pressure, establishing an inverse relationship between CSF flow and neuronal activity, as shown in young individuals^18,22^. Based on this, we predicted that CSF inflow would decrease with the onset of visual stimulation and increase after its offset in healthy older adults as well. Our findings confirmed this prediction. In these healthy individuals, we observed a decrease in CSF inflow concurrent with the onset of visual stimulation, with a notable reduction to -1.91% by 7s after stimulus onset. This decrease followed a linear trend across the first 7s [decreasing linear trend of CSF across 1-7s after stimulus onset, F(1,23)=5.525, P=0.028, partial η^2^=0.194; **Fig. 1d**]. The trough at 7s was significantly lower than the baseline CSF inflow [paired t-test, 7s vs 0s post-stimulus onset: T(23)=-2.601, P=0.016]. After this decline, CSF inflow began to rise [increasing linear trend of CSF across 0-15s after stimulus offset, F(1,23)=14.425, P<0.001, partial η^2^=0.385], peaking at 2.18% 10s after stimulus offset (18s after onset). This peak was significantly higher than baseline CSF inflow [paired t-test, 18s vs 0s post-stimulus onset: T(23)=3.153, P=0.004].

Notably, when examining our glaucoma counterparts, our results revealed that this stimulus-locked pattern of CSF inflow became less apparent with increasing severity of glaucoma (**Fig. 1e,f**). In individuals with early glaucoma, CSF inflow initially showed a stimulus-locked decrease [1-7s after the stimulus onset, F(1,22)=10.697, P=0.003, partial η^2^=0.327], followed by an increasing trend [0-15s after the stimulus offset, F(1,22)=4.476, P=0.046, partial η^2^=0.169]. However, this pattern was absent in individuals with advance glaucoma [decreasing linear trend of CSF across 1-7s after the stimulus onset, F(1,27)=1.275, P=0.269, partial η^2^=0.045; increasing linear trend of CSF across 0-15s after the stimulus offset, F(1,27)=0.869, P=0.359, partial η^2^=0.031].

Additionally, in the power spectral density of CSF inflow signals, healthy individuals displayed a distinct peak at the stimulation frequency (0.0417 Hz; see **Fig. 2a,b** for power spectral density of visual cortex BOLD and CSF inflow). However, individuals with both early and advanced glaucoma lacked this peak, suggesting impaired CSF inflow responses to visual stimuli. When no visual stimulation was presented during resting-state fMRI, no such peak was observed, further confirming that the peak at the stimulation frequency was specifically driven by visual stimulation (see **Extended Data** Fig. 2 for power spectral density of visual cortex BOLD and CSF inflow signals during rest).

**Fig. 2:**
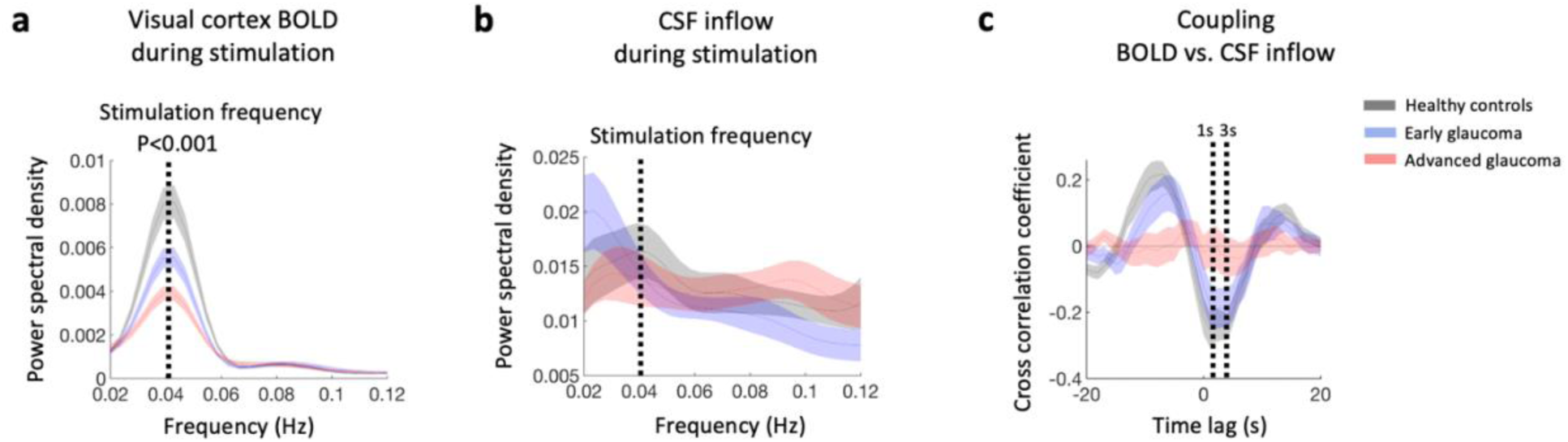
The power spectral density of visual cortex BOLD signals and CSF inflow, and the BOLD-CSF coupling were altered in glaucoma in task-based fMRI. **a**, The power spectral density of the visual cortex BOLD signals indicated that its power at the stimulation frequency (0.0417 Hz, dotted line) declined with increasing severity of glaucoma [main effect of group, F_welch_(2, 42.614)=14.730, P<0.001, partial η^2^=0.319; healthy controls vs. early glaucoma, Games-Howell-corrected P=0.016, 95% confidence interval (CI)=0.0004 to 0.0046; healthy controls vs. advanced glaucoma, Games-Howell-corrected P<0.001, 95% CI=0.0022 to 0.0061; early glaucoma vs. advanced glaucoma, Games-Howell-corrected P=0.012, 95% CI=0.0003 to 0.0030]. **b,** The power spectral density of the CSF inflow indicated that the stimulation-related peak was apparent only in healthy individuals. **c,** The cross-correlation between the BOLD response and CSF inflow showed a maximum anti-correlation at a time lag of 1s (with a low plateau at around 1-3s, dotted lines) in healthy individuals. This correlation coefficient at a lag of 1s was attenuated with increasing glaucoma severity. The data represent mean ± S.E.M. across subjects.

### Coupling between BOLD response and CSF inflow is impaired in individuals with glaucoma

A plausible mechanism for the impaired stimulus-locked pattern of CSF inflow in glaucoma is the reduced neuronal activity triggered by the stimulus. When neuronal activity is compromised, it results in a smaller increase in blood volume^24^. This diminished blood volume may, in turn, weaken its ability to regulate CSF inflow in glaucoma patients. Based on this, we predicted that the coupling between the BOLD response and CSF inflow might be attenuated in glaucoma. To test this prediction, we computed the strength of the coupling between the BOLD response and CSF inflow using cross-correlation analysis.

In healthy individuals, maximal anti-correlation occurred at a lag of 1s (with a low plateau at around 1-3s), indicating that the CSF signal lagged behind the BOLD signal by approximately 1s (**Fig. 2c**). To assess its reliability, we examined the cross-correlation between the modeled BOLD signal, generated from a standard hemodynamic response function and the actual CSF signal obtained from healthy individuals, and observed consistent results (**Extended Data** Fig. 3). Therefore, we selected the correlation coefficient at a lag of 1s as the indicative coupling strength between the BOLD and CSF inflow.

This coupling was weakest in advanced glaucoma [main effect of group, F(2,72)=4.738, P=0.012, partial η^2^=0.116; healthy controls vs. early glaucoma, Bonferroni-corrected P=1.000, 95% CI=- 0.287 to 0.132; healthy controls vs. advanced glaucoma, Bonferroni-corrected P=0.011, 95% CI=- 0.443 to -0.044; early glaucoma vs. advanced glaucoma, Bonferroni-corrected P=0.142, 95% CI=- 0.368 to 0.036]. Furthermore, this measure of coupling strength correlated with some clinical ophthalmic measures that reflected the severity of glaucoma [visual field MD: r=-0.327, P=0.006; foveal threshold: r=-0.467, P<0.001; cup-to-disc ratio (C/D): r=0.330, P=0.005; macular ganglion cell-inner plexiform layer (mGCIPL) thickness: r=-0.247, P=0.047; neuroretinal rim (NRR) area: r=-0.288, P=0.014; retinal structure index: r=-0.289, P=0.021; **Extended Data** Fig. 4]. These results suggest that the coupling between the BOLD signal and CSF inflow deteriorates with worsening glaucoma.

For complementary analysis, we also computed another measure of coupling^21,22^, derived from the cross-correlation between the negative derivative of the BOLD signal (representing contraction of blood volume) and CSF inflow. The results also showed compromised coupling between the two (see **Extended Data** Fig. 5).

Diminished magnitude of the late CSF surging signal in glaucoma can be attributed to a flatter slope in the early BOLD response, with this relationship mediated by a shallower CSF trough.

In addition to the disrupted coupling between the BOLD response and CSF inflow, we predicted that the impaired CSF dynamics in glaucoma could be linked to an impaired BOLD response. Specifically, we hypothesized that the BOLD response would contribute to subsequent CSF signals over time. For instance, the early rising BOLD slope (4-7s post-stimulus onset) would influence the CSF trough (7s post-stimulus onset), which would then affect the CSF surge and its subsequent lingering signal (8-15s post-stimulus offset). Similarly, the early BOLD slope (4-7s post-stimulus onset) could affect the BOLD peak (11-12s post-stimulus onset), which in turn would influence the CSF surge and the following lingering signal (8-15s post-stimulus offset). To investigate these spatiotemporal relationships, we examined the directional influences between these factors using a mediation model.

The late CSF surging signal was first defined as the CSF inflow signal from around its peak until the next stimulus onset. In healthy controls, CSF inflow peaked at about 10s after stimulus offset, while in early and advanced glaucoma patients, the peak occurred at about 8s post-stimulus offset. This peak CSF inflow then stayed until the next stimulus onset, showing no significant decreasing linear trend [no linear trend of CSF; healthy controls, F(1,23)=1.109, P=0.303, partial η^2^=0.046; early glaucoma, F(1,22)=2.057, P=0.166, partial η^2^=0.086; advanced glaucoma, F(1,27)=0.063, P=0.804, partial η^2^=0.002]. To quantify this late CSF surging signal for group comparisons, we calculated the average CSF inflow from 8s after stimulus offset until just before the next stimulus onset (8-15s post-stimulus offset; **Fig. 3a**). This late CSF signal significantly decreased with more severe glaucoma [main effect of group, F(2,72)=4.555, P=0.014, partial η^2^=0.112; healthy controls vs. early glaucoma, Bonferroni-corrected P=0.636, 95% CI=-0.405 to 1.262; healthy controls vs. advanced glaucoma, Bonferroni-corrected P=0.011, 95% CI=0.178 to 1.768; early glaucoma vs. advanced glaucoma, Bonferroni-corrected P=0.304, 95% CI=-0.259 to 1.349; **Fig. 3b**] and correlated with some clinical ophthalmic measures representing the severity of glaucoma [visual field MD: r=0.262, P=0.028, foveal threshold: r=0.269, P=0.027; C/D: r=-0. 298, P=0.011; NRR area: r=0.322, P=0.006; retinal structure index representing the integrity of retina structure (see **Methods** for details): r=0.266, P=0.034; **Extended Data** Fig. 6]. These results suggest that individuals with glaucoma experience a diminishing late CSF surging signal, which worsens with disease severity. When using a wider time window for the average CSF inflow from 0s after stimulus offset until the next stimulus onset as an alternative to the average CSF inflow starting from 8s after stimulus offset, we observed a consistent reduction in the late CSF surging signal with increasing glaucoma severity [main effect of group, F(2,72)=3.371, P=0.040, partial η^2^=0.086].

**Fig. 3:**
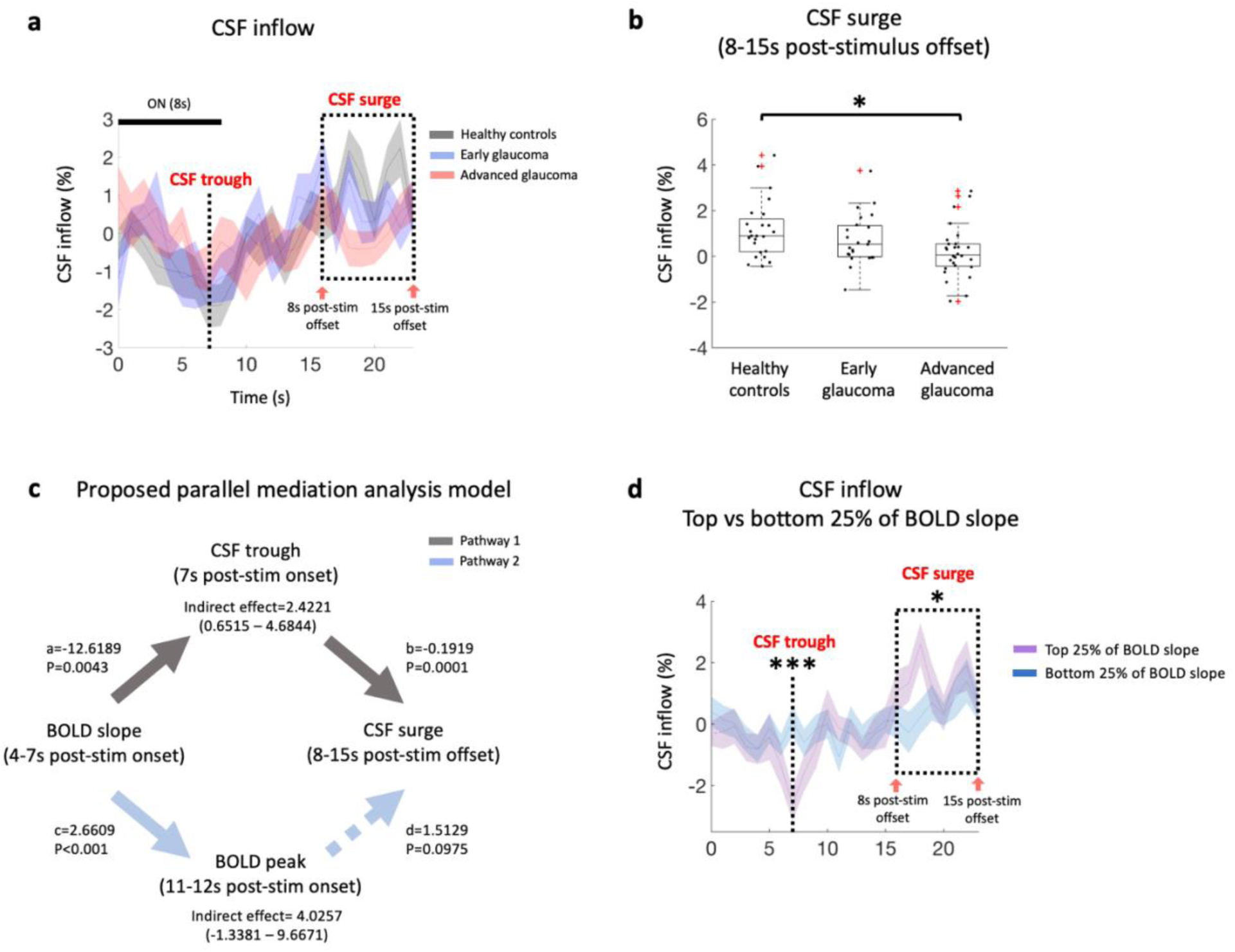
The reduction in the late stimulation-induced CSF surge was attributable to a flatter early BOLD slope, with the CSF trough serving as a mediator. **a**, The time courses of CSF inflow across the three subject groups (healthy individuals, early glaucoma, and advanced glaucoma) showed that the most noticeable differences between groups occurred between 8 and 15s post-stimulus offset (i.e., 16-23s post-stimulus onset; dashed box). **b,** CSF surge during this 8-15s post-stimulus offset decreased as glaucoma severity increased. The data are represented using box and whisker plots, with outliers plotted as red plus signs. **c,** The proposed parallel mediation model links the early BOLD slope, BOLD peak, CSF trough, and CSF surge. The parallel mediation analysis revealed that the early BOLD slope influences CSF surge through the CSF trough, but not through the BOLD peak. **d,** Individuals with the top 25% of BOLD slope exhibited significantly deeper CSF troughs and higher CSF surges compared to those with the bottom 25% of BOLD slope. In a and d, the data represent mean ± S.E.M. across subjects. *Bonferroni-corrected P<0.05. ***Bonferroni-corrected P<0.001.

Given the reduction in the late CSF rebounding signal observed in glaucoma, we tested two possible pathways that could mediate this process (**Fig. 3c)**. First, a flatter early BOLD slope (4-7s post-stimulus onset) could lead to a weaker suppression of CSF inflow, resulting in a shallower CSF trough (7s post-stimulus onset). This, in turn, could limit the late CSF surging signal (8-15s post-stimulus offset). Second, a flatter early BOLD slope (4-7s post-stimulus onset) could result in a reduced BOLD peak (11-12s post-stimulus onset), which again contributes to a decrease in the late CSF surging signal (8-15s post-stimulus offset). In essence, the first pathway proposed that the relationship between the early rising BOLD slope and the late CSF surge is influenced by the CSF trough, while the second pathway proposed an influence of such a relationship through the BOLD peak. To investigate these directional influences, we conducted a parallel mediation analysis. The results revealed that the early BOLD slope influences the late CSF surge through the CSF trough (indirect effect=2.4221, standard error of mean (SEM)=1.0195, 95% CI =0.6515 to 4.6844), but not through the BOLD peak (indirect effect=4.0257, SEM=2.8245, 95% CI= - 1.3381 to 9.6671). The direct effect of the BOLD slope on the CSF surge was not significant (direct effect=-1.9761, SEM=2.9904, P=0.5109), indicating an indirect relationship between these two variables. This observation suggests that the influence of the early BOLD slope on the late CSF surging signal is mediated through the CSF trough. Simple correlations between these factors are illustrated in **Extended Data** Fig. 7. To visually contrast the impact of the early BOLD slope on the subsequent CSF inflow signals, we categorized individuals into high-slope (top 25%) and low-slope (bottom 25%) groups based on their BOLD response and examined their CSF trough and surge properties (**Fig. 3d)**. Consistent with the hypothesis, individuals with steeper BOLD slopes exhibited deeper CSF troughs (F(1,34)=14.392, P<0.001, partial η^2^=0.297) and higher late CSF surge (F(1,34)=4.430, P=0.043, partial η^2^=0.115) compared to those with flatter BOLD slopes.

## Discussion

In this study, we unveiled a compromise in the dynamics of CSF inflow triggered by visual stimulation in glaucoma, an age-related neurodegenerative disease of the visual system. Our findings indicate that as glaucoma severity increases, the distinct patterns of both the early stimulus-induced decreases in CSF inflow and the late increases in CSF inflow locked to the stimulus offset become less pronounced. This disruption is characterized by a weakened coupling between the BOLD response and CSF inflow. Furthermore, we observed that this compromised CSF dynamics can be attributed to an impaired BOLD response. Specifically, our mediation model suggests that a flatter slope of the visual cortex BOLD response leads to a shallower CSF trough, which in turn diminishes the subsequent CSF surging signal. This indicates that impairments in the early BOLD response contribute to the subsequent CSF signals. Our results provide compelling evidence that the visually driven CSF inflow towards the fourth ventricle is compromised in glaucoma due to an impaired early BOLD response, suggesting that alterations in CSF flow extend beyond the optic nerve into the intracranial system.

Our results of the reduced early BOLD slope in the visual cortex and flatter CSF patterns are likely due to the impaired neuronal activity in glaucoma patients. Under normal physiological conditions, neuronal activity triggers arteriolar dilation, leading to increased blood volume through neurovascular coupling^24^. This vascular response, demonstrated by an increase in the BOLD signal, prompts compensatory inverse adjustments in CSF inflow^18^. Previous glaucoma research has demonstrated that the neuronal activity in the primary visual cortex is both reduced^19^ and delayed^20^ in response to visual stimulation. Additionally, fMRI study has shown that BOLD responses are proportional to neuronal firing rates, with a 1% change in the fMRI signal corresponding to approximately 9 spikes per second per neuron^25^. Thus, the impaired neuronal firing rates in glaucoma likely result in a less pronounced increase in blood volume, producing a flatter early BOLD slope in the visual cortex observed in this study. As the BOLD slope flattens, its suppressive effect on CSF inflow weakens, leading to an elevated CSF trough. This elevated CSF trough, in turn, limits the subsequent rebound of CSF inflow. Taken together, these findings suggest that BOLD impairments in the visual cortex, due to diminished neuronal activity, contribute to the reduced CSF surge, with the CSF trough acting as a mediator.

While impaired neuronal activity is considered the major contributor to the flatter BOLD slope in the visual cortex of glaucoma patients, other factors, such as disruptions in neurovascular coupling, vascular reactivity, and modulators of neurovascular mechanisms, cannot be excluded. For example, if blood vessels are unable to respond properly to neuronal activity, this could result in a flatter BOLD slope, even if neuronal activity remains intact. Indeed, impaired neurovascular coupling has been reported in the visual cortex of glaucoma patients^26,27^. Similarly, alterations in the blood vessels’ response to metabolic demands, hormones, and blood pressure could also affect the BOLD signal^28^. Additionally, changes in factors that modulate neurovascular mechanisms, such as cholinergic basal forebrain neurons^29^, might influence top-down processes, thereby affecting the visual cortex BOLD signal. At this stage, the relative contributions of these factors to the flattened BOLD slope are not fully understood. It is possible that a combination of these impairments contributes to the observed changes in the BOLD slope in glaucoma. Further research is warranted to disentangle these contributions.

A crucial implication of the reduced CSF surging signal is its potential to hinder the clearance of neurotoxic waste products. Earlier research suggests that vasomotion, a rhythmic vessel movement, may act as the mechanism linking neuronal activity, CSF flow, and metabolic waste clearance^30^. Arteries naturally undergo vasomotor oscillations at around 0.1 Hz, causing fluctuations in arteriole diameter and subsequent changes in brain tissue oxygenation^31^. When neurons are stimulated, arteriolar dilation occurs^32^, and the ultra-slow fluctuations of neuronal activity synchronize with changes in nearby arteriole diameter^33^, facilitating the clearance process. Studies have shown that visually evoked vasomotion enhances clearance rates^30^, and visually evoked gamma oscillations result in a reduction in amyloid β and tau^34^. Therefore, a decrease in the visually driven CSF surge could potentially impede the clearance process. If this is the case, glaucoma patients could experience a series of less effective visually driven CSF surge and impaired brain waste clearance during vision-related daily activities. Supporting this, research indicates that glaucoma is associated with the accumulation of toxic waste products along the visual pathway, from the retina^9,10^, to the primary visual cortex^11^. Additionally, a reduction in GABA, which is vulnerable to toxic metabolites^15,16^ has been observed in the visual cortex of glaucoma patients^1^. A recent study suggests that GABA promotes the brain waste clearance process^17^, implying that reduced GABA could impair clearance and create a vicious cycle in glaucoma. Moreover, hypertension, a risk factor for primary open-angle glaucoma^35^, was shown to be associated with less efficient perivascular pumping, which drives CSF flow^36^. However, since the current study did not directly measure clearance rates or perivascular pumping, it remains to be seen whether the reduced CSF surge is functionally related to the impaired clearance of neurotoxic waste products.

Our results suggest that alterations in CSF flow may extend beyond the optic nerve into the fourth ventricle, encompassing ocular and other intracranial CSF spaces. This broader impact is plausible given the interconnected nature of these spaces through which CSF circulates. Supporting this, previous studies have observed impaired CSF exchange between the intracranial CSF spaces and the optic nerve subarachnoid space^37^, slowed CSF flow in the optic nerve subarachnoid space^6,7^, and impaired CSF entry into the optic nerve^38,39^ in glaucoma. Conversely, some studies have reported an opposite trend of enhanced entry into the optic nerve^5,40^. Therefore, future studies should broaden their investigation to include other ocular and intracranial CSF spaces. Additionally, since body position, whether upright or supine, is thought to affect CSF flow^41,42^, further research is needed to disentangle the effects of body position on CSF dynamics. Other considerations include the impact of sleep on CSF flow in glaucoma, as CSF flow^22^ and the clearance process^43-45^ are most active during sleep, and poor sleep quality is associated with glaucoma^46^. Future research should evaluate CSF dynamics during different sleep states in glaucoma patients.

To conclude, our study reveals that the dynamics of CSF inflow driven by visual stimulation is impaired in the intracranial system of glaucoma patients, potentially due to impaired neuronal responses. Our results suggest that glaucoma has a broader impact on CSF dynamics, potentially affecting the intracranial CSF spaces and neural tissues beyond the optic nerve in a systemic manner. Additionally, our non-invasive approach highlights a novel method for monitoring CSF flow in glaucoma patients and paves the way for a better understanding of glaucoma pathogenesis, as well as the development of innovative treatments to reduce the prevalence of this world’s leading cause of irreversible but preventive age-related blinding disease.

## Methods

### Subjects

Twenty-three early glaucoma patients (age = 65.17 ± 1.59 years (mean ± SEM); 30.4% male), twenty-nine advanced glaucoma patients (age = 66.14 ± 1.33 years (mean ± SEM); 55.2% male), and twenty-seven healthy controls (age = 63.93 ± 1.47 years (mean ± SEM); 44.4% male) participated in the study between August 2018 and January 2024 at New York University Langone Health’s Department of Ophthalmology (see **Table 1**. for demographic and clinical information). All participants had a best corrected visual acuity of 20/60 or better and had no present indications of neurological disorders. Individuals were either diagnosed with primary glaucoma, or classified as healthy with no clinical indications of glaucoma. Individuals with any additional retinal disorders were excluded from participation in the study. This study was approved by the Institutional Review Board of New York University Grossman School of Medicine. Before participation, written informed consent was obtained from all subjects.

### Clinical ophthalmic exams

We collected measures of pRNFL thickness, mGCIPL thickness, optic nerve head C/D, and NRR area from all individuals, utilizing the Cirrus spectral-domain OCT device (Zeiss, Dublin, CA, USA). Additionally, we collected functional data from visual field MD and foveal threshold using the Humphrey Swedish Interactive Thresholding Algorithm (SITA) 24-2 standard (Zeiss, Dublin, CA, USA). Individuals with glaucoma were then assigned into early or advanced stage based on the average MD of the left and right eyes. Participants with an average MD greater than -6.0dB were categorized as being in the early stage, while those with an average MD lower than -6.0dB were classified as being in the advanced stage^1,23^.

### MRI data acquisition

Subjects were scanned inside a 3-Tesla MR Prisma scanner (Siemens, Germany) equipped with a 20-channel head coil at the Center for Biomedical Imaging, NYU Langone health, New York University. High-resolution anatomical images were obtained using a multi-echo magnetization-prepared rapid gradient echo sequence (256 slices, voxel size=0.8×0.8×0.8mm^3^, repetition time (TR)=2400ms, echo time (TE)=2.24ms, flip angle=8°, field of view=256mm, bandwidth=210 Hz per pixel). Subsequently, resting-state and task-based functional MR images were acquired using a gradient-echo echo-planar imaging (EPI) sequence (voxel size=2.3×2.3×2.3mm^3^, TR=1000ms, TE=32.60ms). During the resting-state fMRI scan (1 run, scanning duration=480s), subjects were instructed to close their eyes while remaining awake. In the task-based fMRI session (2 runs, scanning duration for 1 run=300s), subjects performed a fixation task while a flickering checkerboard stimulus was presented on the entire screen. In each run, subjects covered one eye at a time with a plastic eye occluder, ensuring that only one eye was stimulated. Four individuals were unable to complete the task-based fMRI session due to technical issues.

### Fixation task

During the fixation task, subjects were instructed to keep their gaze fixed on the center of the screen and press a button whenever the central dot changed color from white to red. The rest of the screen displayed a flickering checkerboard pattern. To avoid scanner instability, we decided to exclude the first and last 6s periods, during which no checkerboard was presented. After the initial 6s period, a 16s “Off” condition was introduced, followed by an 8s “On” condition. Throughout the 300s fixation task, there were 12 “ON” conditions and 11 “OFF” conditions, alternating in order. Each trial was defined as an 8s “On” condition followed by a 16s “Off” condition. In the analysis, the final “ON” condition was excluded because it was not followed by an “OFF” condition, making it impossible to detect changes in CSF signals. As a result, 22 trials across two runs were included in the analysis. The average accuracy of the fixation task (± S.E.M.) was 96.48 ± 0.506%, with no significant differences observed between groups (main effect of group, F_welch_(2, 28.371)=1.076, P=0.355, partial η^2^=0.024).

### Analysis of time-course fMRI data

Task-based and resting-state fMRI data were analyzed using FreeSurfer software version 7.2 (http://surfer.nmr.mgh.harvard.edu/) and MATLAB version R2021a. Preprocessing of the fMRI data involved motion correction without spatial or temporal smoothing, followed by registration to each individual’s structural template. To ensure signal stability, the initial 6 volumes were removed, and voxels exhibiting spikes greater than 10 standard deviations from the mean were excluded. Subsequently, linear and quadratic trends were removed from the time series of each voxel, and the BOLD signal was converted to percent signal change by dividing it by the mean value of the time series. Then we extracted the mean BOLD signal from the visual cortex mask (hOc1-hOc4v) generated from FreeSurfer’s cortical reconstruction process and labeled it as visual cortex BOLD. We also extracted the mean BOLD signal from the entire gray matter mask and named it as the global BOLD signal. For the CSF inflow signal, we selected voxels that spatially overlap with the fourth ventricle from the bottom slice of the fMRI volume and extracted the mean BOLD signal from this mask. For each individual, the time courses of the BOLD and CSF signals were averaged across 22 trials. Cross-correlation analysis was then performed using MATLAB’s ‘xcorr’ function, over time lags ranging from -20s to 20s.

### Analysis of power spectral density of fMRI data

For the power spectral density analysis, the preprocessed fMRI data were linearly detrended, bandpass filtered between 0.01-0.15 Hz and normalized using AFNI (version AFNI_23.3.04) function ‘3dTproject’. Then we computed the power spectral density using MATLAB function ‘pwelch’ with default parameters, which divided the dataset into eight equal-length segments with 50% overlap, applying Hamming window.

### Statistics and reproducibility

We conducted parametric tests with a significance level of P<0.05 for all statistical comparisons. The homogeneity of variances was assessed using the Levene’s test, and in cases of violation, we employed the Welch test. Bonferroni corrections were applied for post-hoc tests. Principal component analysis was used to extract a common component, termed the “retinal structure index” from the retinal variables, including pRNFL thickness, mGCIPL thickness, optic nerve head C/D, and NRR area, with an eigenvalue criterion set above one. The sample size was not predetermined but was comparable to or even larger than those reported in previous studies^47,48^.

### Apparatus

Visual stimuli were generated using MATLAB along with Psychophysics Toolbox 3. These stimuli were presented using an MRI-compatible projector with a resolution of 1024×768 pixels and a refresh rate of 60 Hz.

## Data availability

The data supporting the results and figures are available at: https://osf.io/myuek/

## Code availability

The scripts used for analysis are available at: https://osf.io/myuek/

## Acknowledgements

The authors express gratitude to Ms Tonya Robins, Zena Moore, Jamika Singleton-Garvin, and members of the Neuroimaging and Visual Science Laboratory at New York University Grossman School of Medicine for their assistance with subject recruitment and technical support. This work is supported in part by the National Institutes of Health R01-EY028125, and P41-EB017183 (Bethesda, Maryland), BrightFocus Foundation G2019103, and G2021001F (Clarksburg, Maryland), and an unrestricted grant from Research to Prevent Blindness to NYU Langone Health Department of Ophthalmology (New York, New York).

## Supplementary information

**Extended Data Fig. 1:**
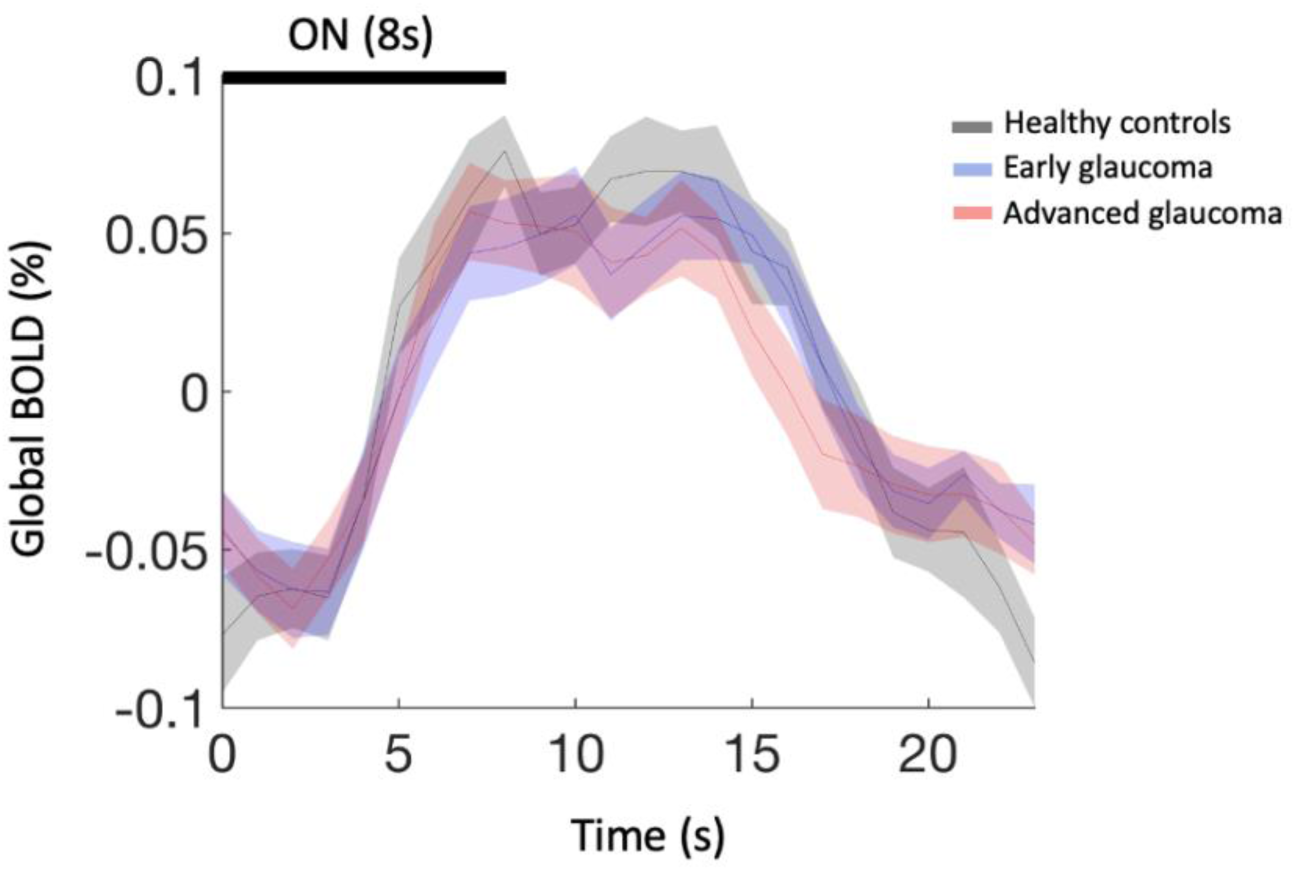
Time courses of global BOLD signals in healthy controls, individuals with early glaucoma, and individuals with advanced glaucoma during task-based fMRI. The average BOLD amplitude (7-14s post-stimulus onset) was comparable across groups (main effect of group, F(2, 72)=0.700, P=0.500, partial η^2^=0.019). The data represent mean ± S.E.M. across subjects.

**Extended Data Fig. 2:**
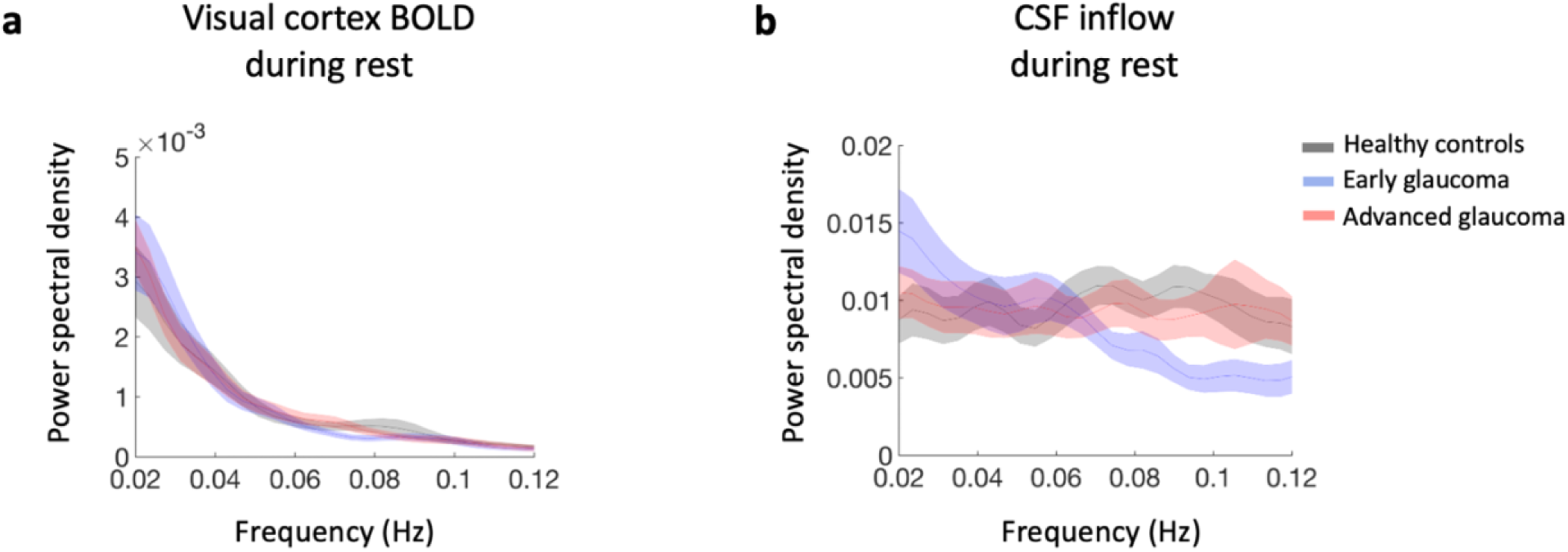
Power spectral density of visual cortex BOLD signals and CSF inflow during an 8-min resting-state fMRI scan without a visual task. Unlike the power spectral density observed in task-based fMRI scan, there was no noticeable peak around 0.0417 Hz in the power spectral density of **a,** visual cortex BOLD or **b,** CSF inflow during the resting-state fMRI scan of healthy controls. The data represent mean ± S.E.M. across subjects.

**Extended Data Fig. 3:**
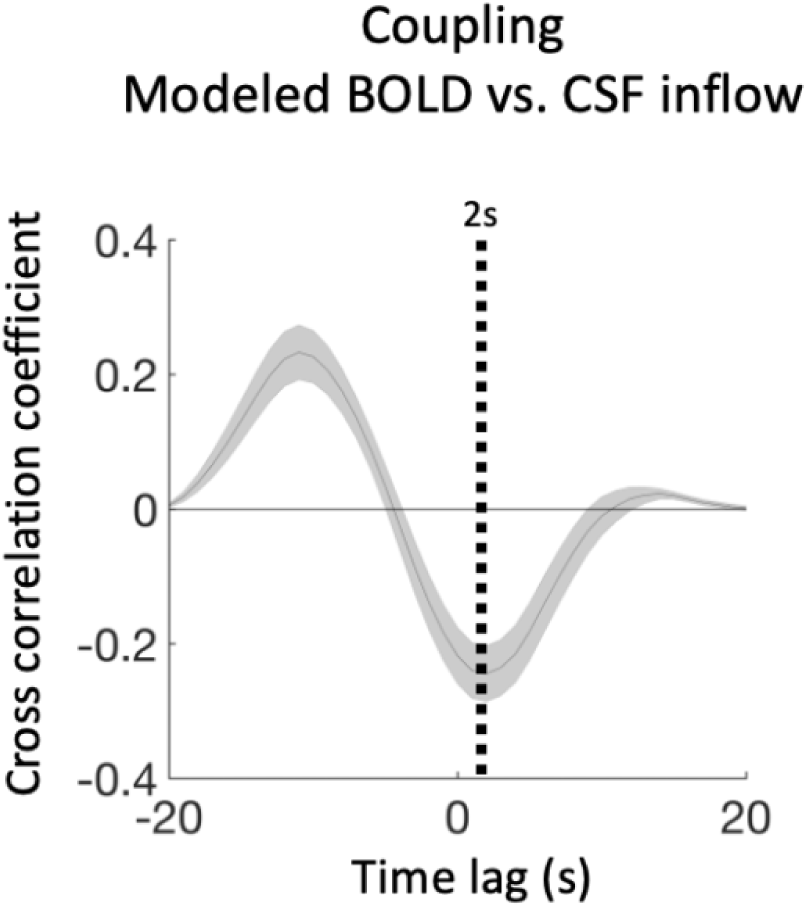
Cross-correlation between the modeled BOLD signal and actual CSF inflow recorded during task-based fMRI from healthy controls. The cross-correlation of the modeled BOLD showed a maximum anti-correlation with the actual CSF inflow at a time lag of 2s. The data represent mean ± S.E.M. across subjects.

**Extended Data Fig. 4:**
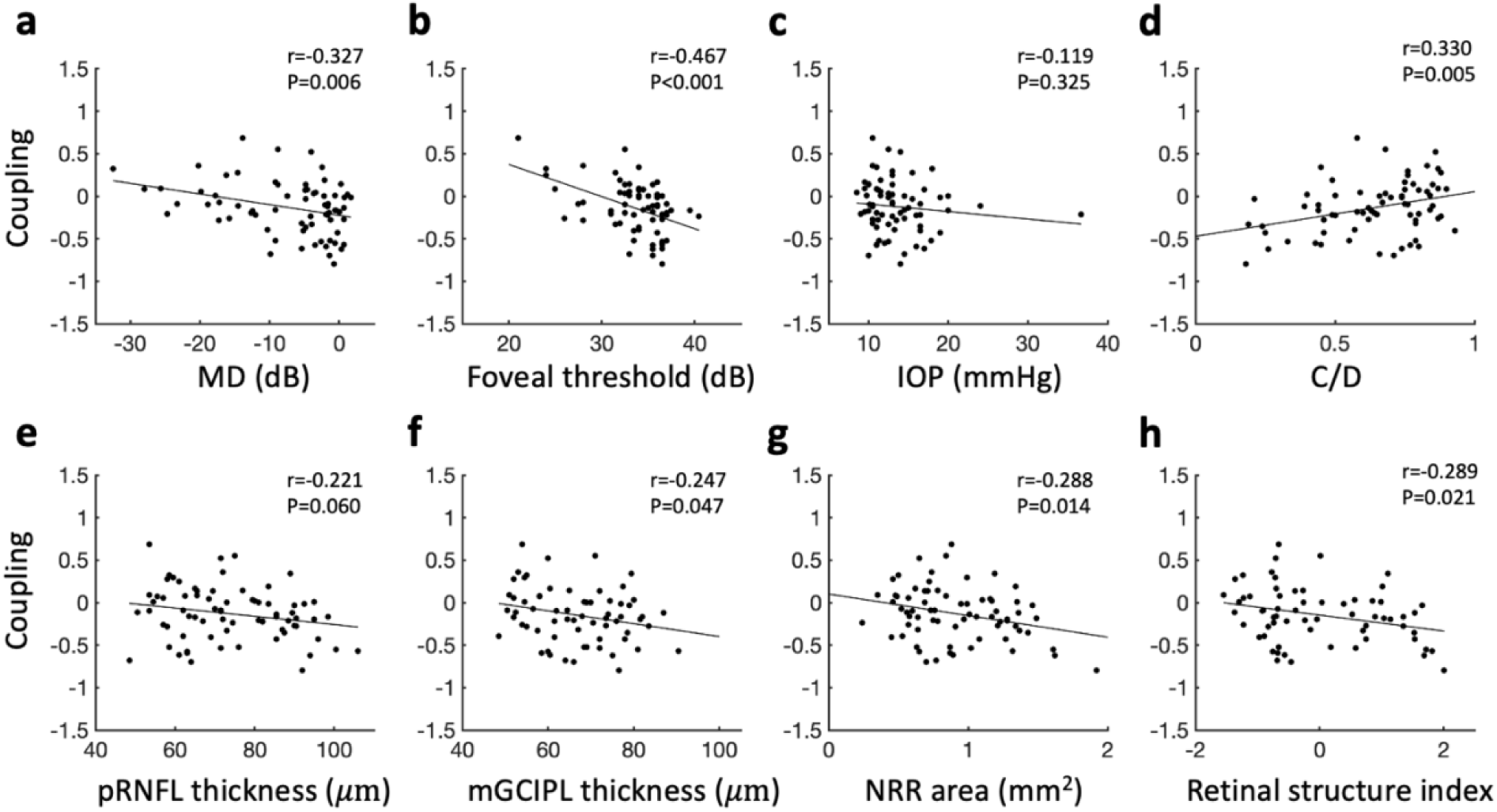
Coupling between task-based visual cortex BOLD response and CSF inflow was found to correlate with various clinical ophthalmic measures. The scatterplots show the correlation of coupling strength with the following measures: **a,** visual field mean deviation (MD), **b,** foveal threshold, **c,** intraocular pressure (IOP), **d,** cup-to-disc ratio (C/D), **e,** peripapillary retinal nerve fiber layer (pRNFL) thickness, **f,** macular ganglion cell-inner plexiform layer (mGCIPL) thickness, **g,** neuroretinal rim (NRR) area, and **h,** retinal structure index (derived from principal component analysis of retinal variables, including pRNFL thickness, mGCIPL thickness, optic nerve head C/D, and NRR area). Significant correlations were observed with MD, foveal threshold, C/D, mGCIPL thickness, NRR area, and the retinal structure index.

**Extended Data Fig. 5:**
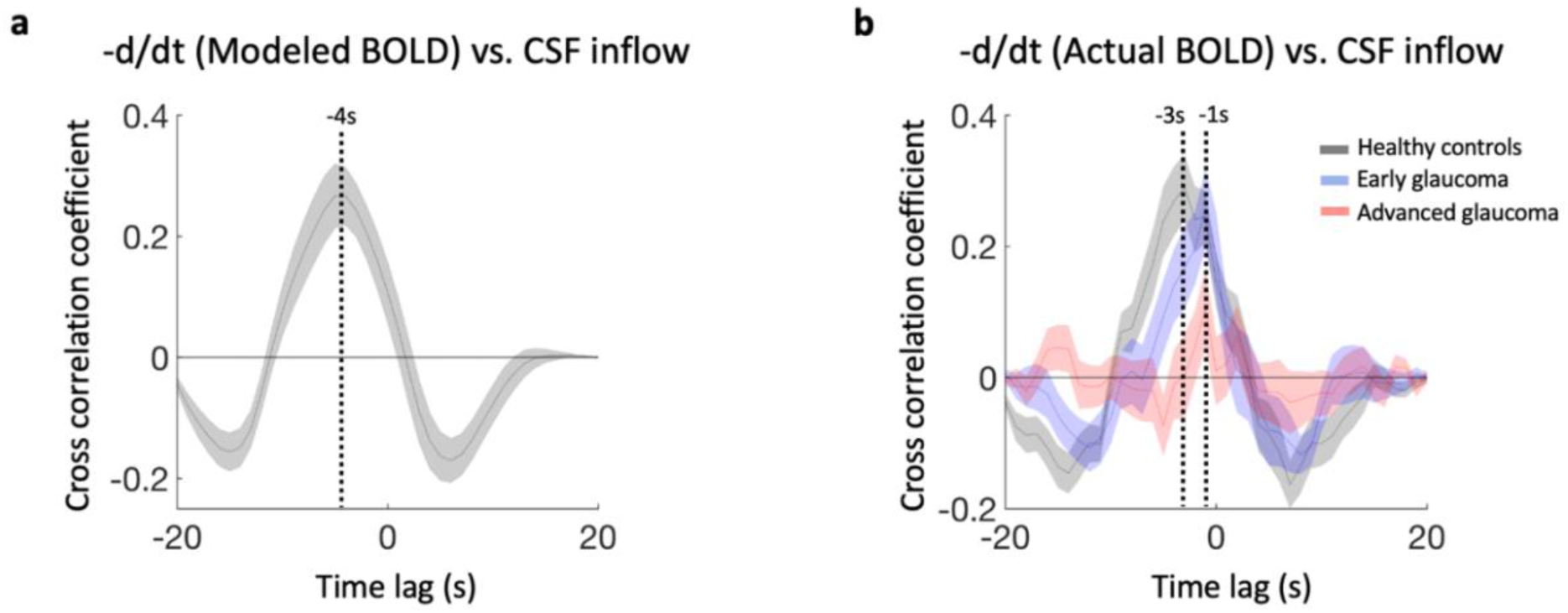
Cross-correlations between the negative derivatives of the BOLD signals and CSF inflow in task-based fMRI. **a**, The maximum correlation was observed at a lag of approximately -4s when using modeled BOLD signals and **b,** at a lag of -3s when using actual visual cortex BOLD from healthy individuals. The correlation coefficient at a lag of -3s diminished with increasing glaucoma severity (main effect of group, F(2,72)=6.087, P=0.004, partial η^2^=0.145; healthy controls vs. early glaucoma, Bonferroni-corrected P=0.488, 95% CI=-0.091 to 0.337; healthy controls vs. advanced glaucoma, Bonferroni-corrected P=0.003, 95% CI=0.084 to 0.491; early glaucoma vs. advanced glaucoma, Bonferroni-corrected P=0.162, 95% CI=-0.041 to 0.371). Additionally, the time lag for maximum correlation shifted to -1s in both stages of glaucoma. The data represent mean ± S.E.M. across subjects.

**Extended Data Fig. 6:**
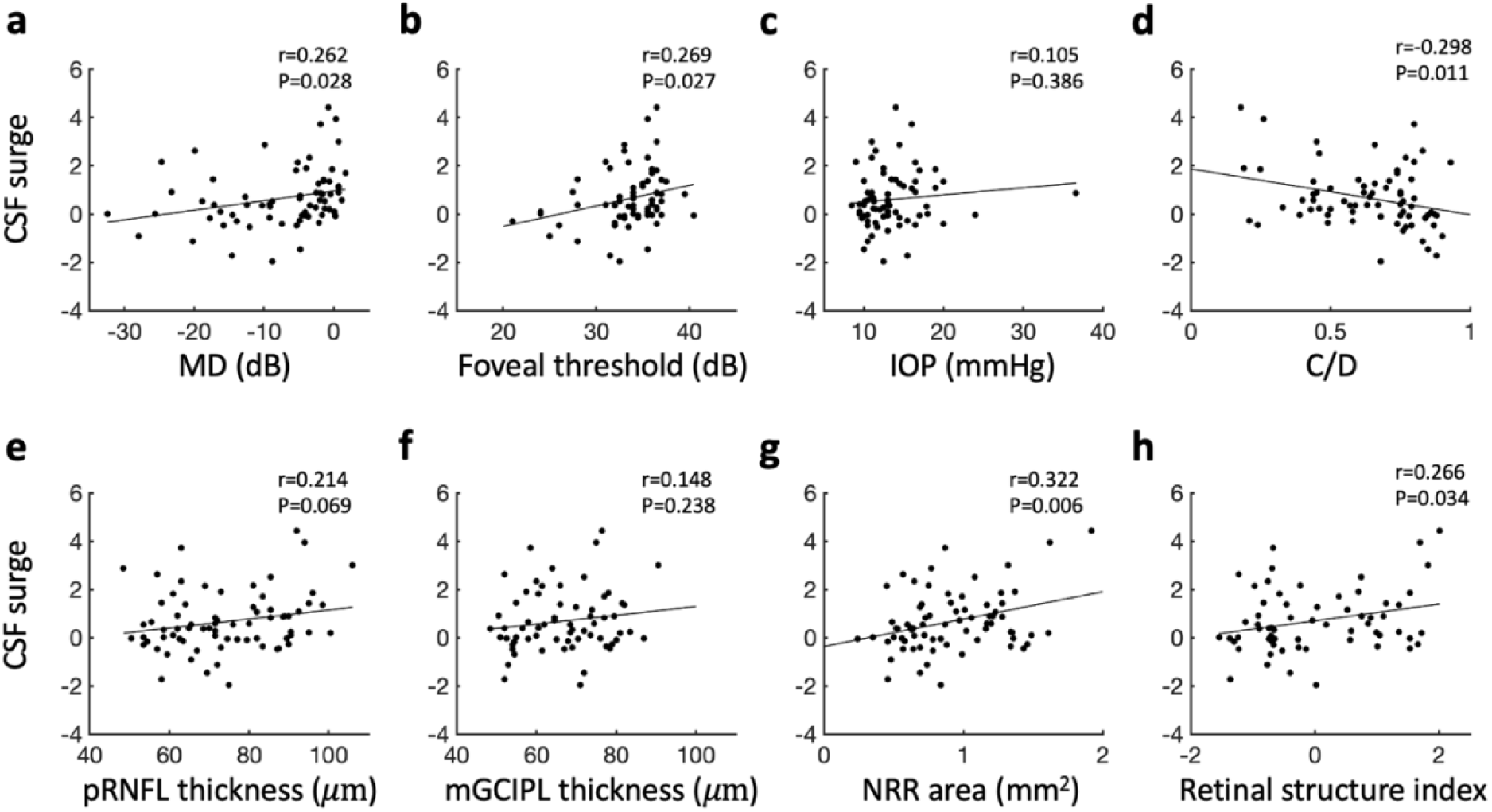
CSF surge observed during task-based fMRI was correlated with various clinical ophthalmic measures. The scatterplots illustrate the correlation between CSF surge and the following measures: **a,** visual field mean deviation (MD), **b,** foveal threshold, **c,** intraocular pressure (IOP), **d,** cup-to-disc ratio (C/D), **e,** peripapillary retinal nerve fiber layer (pRNFL) thickness, **f,** macular ganglion cell-inner plexiform layer (mGCIPL) thickness, **g,** neuroretinal rim (NRR) area, and **h,** retinal structure index (derived from principal component analysis of retinal variables, including pRNFL thickness, mGCIPL thickness, optic nerve head C/D, and NRR area). Significant correlations were found with MD, foveal threshold, C/D, NRR area, and the retinal structure index.

**Extended Data Fig. 7:**
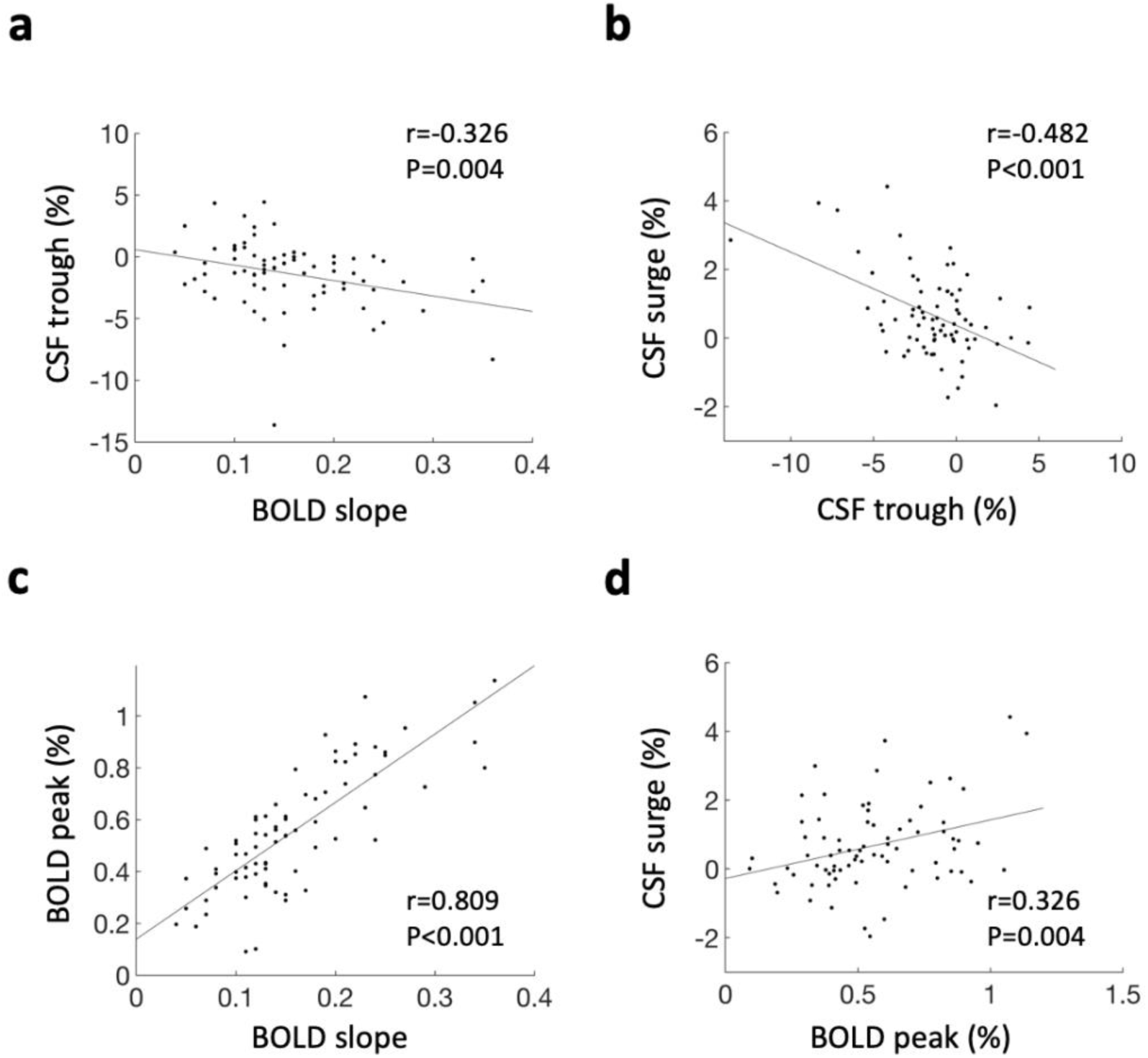
Simple linear correlations between early BOLD slope, BOLD peak, CSF trough, and CSF surge during task-based fMRI. These variables were incorporated into the proposed parallel mediation model. Scatter plots illustrate the relationships between **a,** BOLD slope and CSF trough, **b,** CSF trough and CSF surge, **c,** BOLD slope and BOLD peak, and **d,** BOLD peak and CSF surge.

